# Prediction of Choice From Competing Mechanosensory and Choice-Memory Cues During Active Tactile Decision Making

**DOI:** 10.1101/400358

**Authors:** Campagner Dario, Evans H Mathew, Chlebikova Katarina, Colins-Rodriguez Andrea, Loft S E Michaela, Fox Sarah, Pettifer David, Humphries D Mark, Svoboda Karel, Petersen S Rasmus

## Abstract

Perceptual decision making is an active process where animals move their sense organs to extract task-relevant information. To investigate how the brain translates sensory input into decisions during active sensation, we developed a mouse active touch task where the mechanosensory input can be precisely measured and that challenges animals to use multiple mechanosensory cues. Mice were trained to localise a pole using a single whisker and to report their decision by selecting one of three choices. Using high-speed imaging and machine vision we estimated whisker-object mechanical forces at millisecond resolution. Mice solved the task by a sensory-motor strategy where both the strength and direction of whisker bending were informative cues to pole location. We found competing influences of immediate sensory input and choice memory on mouse choice. On correct trials, choice could be predicted from the direction and strength of whisker bending, but not from previous choice. In contrast, on error trials, choice could be predicted from previous choice but not from whisker bending. This study shows that animal choices during active tactile decision making can be predicted from mechanosenory and choice-memory signals; and provides a new task, well-suited for future study of the neural basis of active perceptual decisions.

**SIGNIFICANCE STATEMENT:** Due to the difficulty of measuring the sensory input to moving sense organs, active perceptual decision making remains poorly understood. The whisker system provides a way forward since it is now possible to measure the mechanical forces due to whisker-object contact during behaviour. Here we train mice in a novel behavioural task that challenges them to use rich mechanosensory cues, but can be performed using one whisker and enables task-relevant mechanical forces to be precisely estimated. This approach enables rigorous study of how sensory cues translate into action during active, perceptual decision making. Our findings provide new insight into active touch and how sensory/internal signals interact to determine behavioural choices.

## INTRODUCTION

Perceptual decision making (Romo and Salinas, 2003; Cohen and Newsome, 2004; Gold and Shadlen, 2007; Carandini and Churchland, 2013; Diamond and Arabzadeh, 2013; Svoboda and Li, 2018) is an active process where a movement of the sense organs – e.g., eyes, ears, nose, fingers or whiskers – is crucial to extract task-relevant information (Gibson, 1962; Yarbus, 1967; Youngentob et al., 1987; Jordan et al., 2018). Our understanding of how the brain translates sensory signals into decisions during active sensation has been held back by the experimental difficulty of measuring sensory input to a moving sense organ. However, new approaches developed for the mouse whisker system provide a way forward (O’Connor et al., 2010b, 2013; Hires et al., 2015; Peron et al., 2015a; Yu et al., 2016). Here, we describe a new tactile task for mice that permits precise monitoring of sensory input during active, perceptual decision making, and thereby identifies specific mechanosensory and choice-memory signals that predict the animals’ choices.

Rats and mice explore objects by probing them with back-and-forth movements of their whiskers (‘whisking’, Vincent, 1912; Welker, 1964), and can solve a wide range of tasks in this way (Hutson and Masterton, 1986; Guić-Robles et al., 1989; Carvell and Simons, 1990; Krupa et al., 2001; Polley et al., 2005; Anjum et al., 2006; Knutsen et al., 2006; Mehta et al., 2007; Favaro et al., 2011; Sofroniew et al., 2014; Fassihi et al., 2014a; Bale et al., 2017; Nikbakht et al., 2018; Evans et al., 2018). Contact causes whiskers to bend and the associated torque (‘bending moment’) is a major driver of spikes fired by Primary Whisker Neurons (PWN) located in the Trigeminal Ganglion (Bush et al., 2016; Campagner et al., 2016; Severson et al., 2017, reviewed by Campagner et al., 2017). However, how such mechanosensory cues translate into perceptual decisions is not fully understood.

Recently, high-speed imaging and machine vision methods have been developed, which make it possible to measure whisker-object forces in behaving animals (Birdwell et al., 2007; O’Connor et al., 2010a; Clack et al., 2012; Pammer et al., 2013; reviewed by Campagner et al., 2017). A head-fixed mouse paradigm, where animals are trained to localise a vertical pole with their whiskers, is advantageous. Head-fixation permits whisker movement and whisker shape to be imaged at high spatiotemporal resolution. The curvature of a whisker bending against a vertical pole can be measured, allowing whisker-object contacts, and associated mechanical forces, to be precisely estimated. Previous studies have used two-choice tasks where animals are trained to report anterior-posterior or medial-lateral pole location by licking (O’Connor et al., 2010a; Pammer et al., 2013; Guo et al., 2014b). Mice solve the anterior-posterior task by learning to focus their whisking on one of the pole locations. In this way, the strength and number of touches allows mice to discriminate pole location (O’Connor et al., 2010a, 2010b, 2013). However, it remains unclear how rodents solve active touch tasks under conditions when these elementary cues are insufficient. Here, we developed a novel, three-choice pole localisation task, where the mechanosensory input guiding decision-making can be precisely measured, and that challenges mice to use cues beyond the strength and number of touches. We identified key mechanosensory cues and discovered that these cues, in conjunction with an internal signal (memory of choices on previous trials) allowed mouse choices to be accurately predicted.

## MATERIALS AND METHODS

All experimental protocols described in this section were approved by both United Kingdom Home Office national authorities and institutional ethical review.

### Surgical procedure and water restriction

Mice (C57; males; N=5; 6 weeks at time of implant) were implanted with a titanium head-bar as detailed in (Campagner et al., 2016). After surgery, mice were left to recover for at least 5 days before starting water restriction (1.5 ml water/day). Training began 7-10 days after the start of water restriction.

### Behavioural apparatus

Mice were trained in a dark, sound-proofed enclosure adapted from O’Connor et al. (2010a) and Campagner et al. (2016). Briefly, a head-fixed mouse was placed inside a Perspex tube, from which its head emerged at one end. The stimulus object was a 1.59 mm diameter, vertical metal pole which could be translated parallel to the anterior-posterior axis of the mouse by a linear stepper motor (NA08B30, Zaber, Vancouver, Canada). To allow vertical movement of the pole into and out of range of the whiskers, the pole was mounted on a pneumatic linear slide (SLS-10-30-P-A, Festo, Northampton, UK), powered by compressed air. The apparatus was controlled from MATLAB (The MathWorks, Inc., Natick, Massachusetts, United States) via a real-time processor (RX8, TDT, Alachua, FL). Mouse response was monitored by two lick ports located anterior to the mouth. Licks were detected as described in O’Connor et al., 2010a (Fig. 1 A and B). Each lick port consisted of a metal tube connected to a water reservoir via a computer-controlled solenoid valve (LHDA1233215H, Lee Company). Lick port position was monitored using an infrared camera (N08CX-Sentient) and adjusted using a micromanipulator.

**Figure 1.**
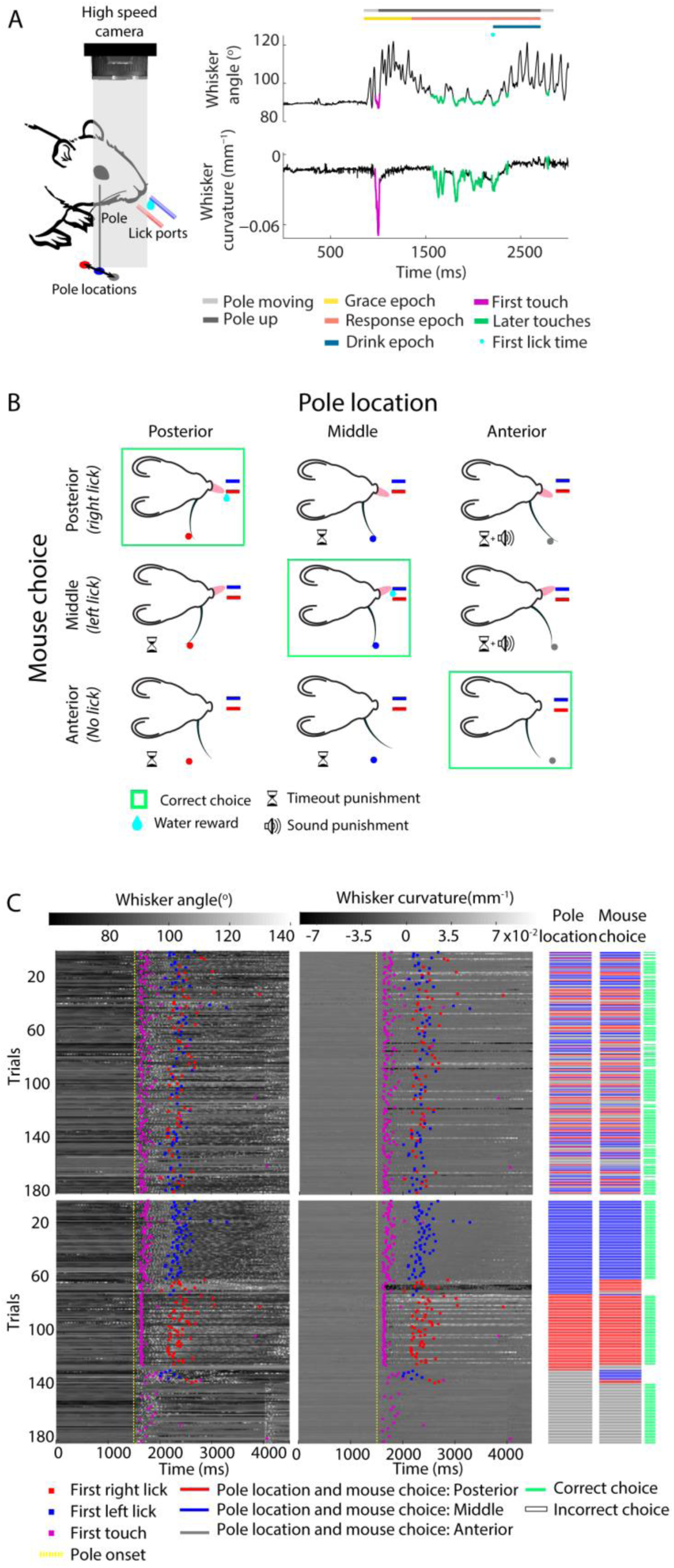
The three-choice object localisation task. **A. Left**: Schematic of the experimental preparation, showing the three pole locations (circles) and the two lick ports. Both lick ports and pole location are colour coded consistently with panel B. Whisker movements and whisker-pole interactions were filmed with a high-speed camera (1000 frame/s). **Right:** Schematic of a correct go trial to illustrate the trial structure (coloured bars, defined in *Materials and Methods*). Whisker angle, whisker curvature and whisker-pole touches were extracted from the high-speed video. Mouse choice was monitored by measuring the time of first lick. **B.** Trial-choice outcomes and how they were rewarded/punished. **C.** Mouse behaviour during an example experimental session. Whisker angle (left) and whisker curvature (right) for each whisker tracked trial (*Material and Methods* and Fig. 3-1 and 3-2). In the top panels, trials are sorted according to chronological order during the session. In the bottom panels, trials are sorted first by pole location, and, within each pole location, by mouse choice.

### Behavioural task

Head-fixed mice were trained to locate a metal pole using their whiskers and to report its position by licking (Fig. 1 B). On each trial, the pole was presented in one of three anterior-posterior locations (posterior, middle and anterior). On trials where the pole was middle or posterior (‘go left location’ or ‘go right location’), the correct response was for the mouse to lick one of the two lick ports. Correct responses were rewarded by a drop of water ∼10μl. In 3 cases (mice 32, 33 and 34), animals were rewarded for licking at the right lick port when the pole was posterior, and for licking at the left lick port when the pole was middle. In 2 other cases (mice 36 and 38), the contingency was reversed. Incorrect responses on go left/right trials (licking the wrong side or not licking at all) were punished by timeout (Fig. 1 B, hourglass symbol). On trials where the pole was anterior (‘no go location’), the correct response was to refrain from licking. Incorrect responses on no go trials (licking) were punished by timeout and tone (frequency 1 kHz; Fig. 1 B, speaker symbol).

#### Trial structure

Each trial started with the pole in its down position, out of reach of the whiskers (Fig 1 A, right panel). Licks during this epoch were ignored. Then, at ‘pole onset’, the pneumatic valve opened, causing the pole to move up within reach of the whiskers (pole travel time ∼0.15 s). As in related previous studies, during training, the sound caused by opening of the valve tended to trigger reflexive licks, unrelated to mouse choice (O’Connor et al., 2010a; Guo et al., 2014b). To exclude these, for a short ‘grace epoch’ following pole onset (typically 0.5 s for the full task, defined below) licks were ignored.

The grace epoch was immediately followed by a ‘response epoch’. During this time period mouse licking could control water delivery (typical duration 2 s for the full task). If, during the response epoch of a go trial, a mouse licked the correct lick port, the first lick triggered the onset of a drink epoch: the water valve opened, making a drop of water available at the lick port. Drink epoch duration varied over the course of training (typically 0.5-2 s). At the end of the ‘drink epoch, the pneumatic valve closed, causing the pole to move back to its down position, and the trial terminated (Fig. 1 A, right panel). If, during the response epoch, a mouse did not lick or licked the incorrect lick port, it was punished by a ‘timeout epoch’ (typically 2-10 s). If, during the response epoch of a no go trial, a mouse did not lick, the trial was terminated at the end of the response epoch, causing the pole to return back to its down position. If, instead, the mouse licked one of the lick ports, there was a timeout epoch, at the end of which the pole returned to its down position.

#### Training protocols

The mouse training process was divided into successive protocols of increasing complexity, following (O’Connor et al., 2010a; Guo et al., 2014b). Transition from one protocol to the next was performed only if the mouse showed stable performance (∼70%) on at least two consecutive days (see Fig. 2 A and Fig. 2-1 A). The typical sequence of training protocols was as follows:

**Figure 2.**
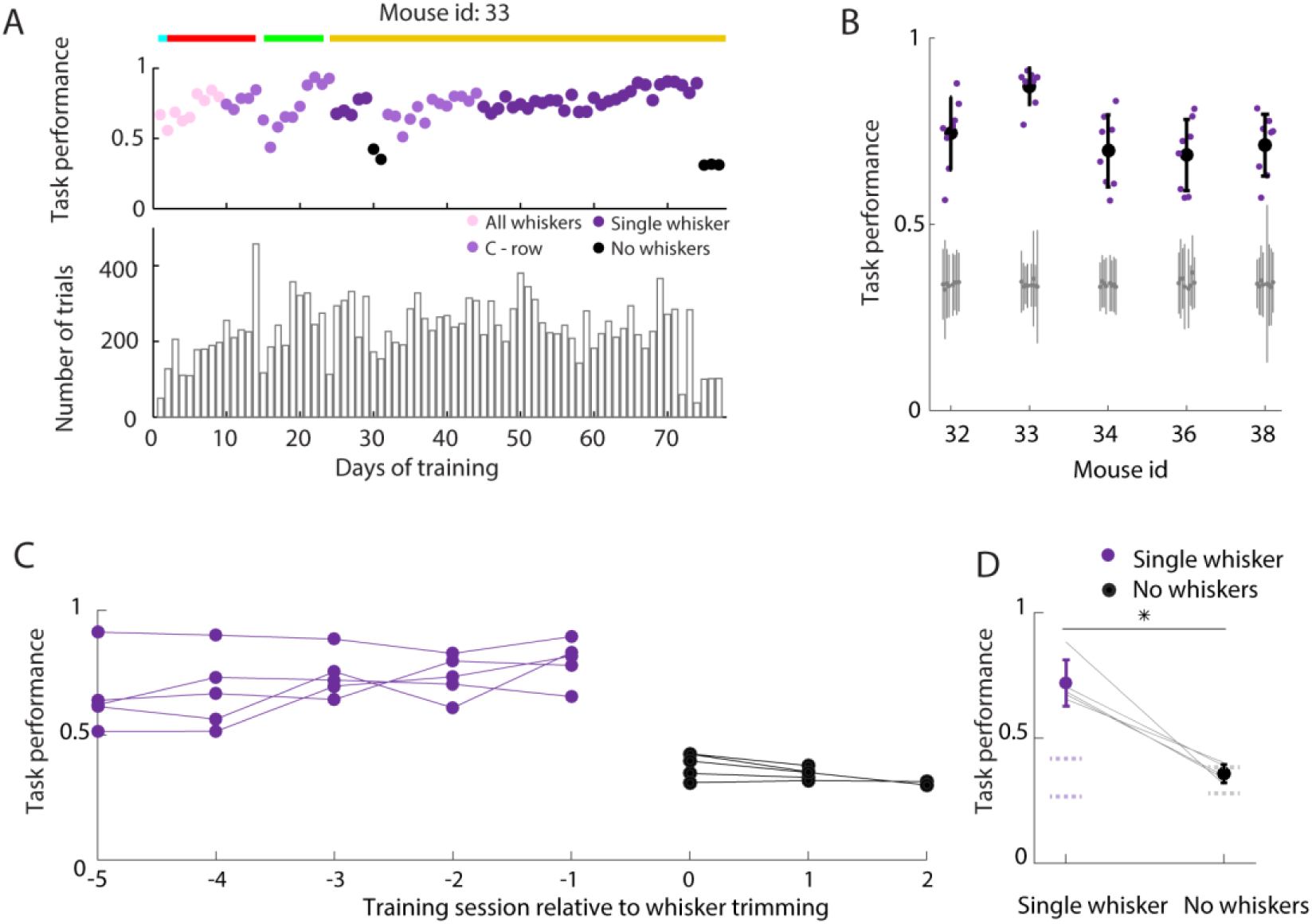
The three-choice object localisation task is whisker-dependent. **A. Top**: Task performance of mouse 33 during the course of training (for analogous data for other mice, see Fig. 2-1 A). The mouse was initially trained with all its whiskers intact. The whiskers were progressively trimmed to one whisker and, finally, as a control, to none. Coloured lines indicate the protocol the mouse was trained on each day: lick (Cyan), go – no go (red), lick left - lick right (green), lick left - lick right – no lick (gold) protocol (protocols detailed in *Materials and Methods*). When cyan and red lines overlap, it indicates that the protocol was switched to go – no go protocol during the same behavioural session. **Bottom**: Total number of trials performed each day. **B.** Stable performance for each mouse during AB trials of the full task with single whisker. Stable sessions selected as detailed in *Materials and Methods*. Purple dots show performance in each session; large black dots and black error bars show mean and SD across selected sessions respectively. Gray dots and gray error bars show chance performance and 95% confidence interval on chance respectively. **C.** Task performance during AB trials in the 5 sessions before (purple) and 2-3 sessions after (black) whisker trimming, for each of 5 whisker trimming tests. **D.** Grand mean task performance on sessions before (dark purple) and after (black) whisking trimming. Error bars: SD .* t-test p = 0.0013. Dotted lines: average chance range (*Materials and Methods*).

##### 1) Lick

First, mice were trained to associate whisker-pole contact with availability of water from the lick ports. Whenever the pole moved up into one of the two go locations, a drop of water was delivered. After a few trials, mice started to lick in response to the pole movement, triggering water delivery via the lick sensor.

##### 2) Go – no go

Next, mice were trained to lick selectively based on pole location. On each trial, the pole was presented in one of two alternative locations: the posterior go location or the anterior no go location. Only one lick port was within reach. The mouse was rewarded for licking when the pole was presented in the go location. The mouse was punished (by timeout) for both false alarms (licking on no go trials) and misses (not licking on go trials). When the mouse reached stable performance (∼70% correct performance) with its full whisker array, all whiskers except for C row were trimmed to fur level. This whisker configuration was maintained by repeated re-trimming over the successive days/weeks. If trimming caused a drop in mouse performance, training continued with a single row of whiskers in the same protocol used before trimming, until performance returned to its pre-trimming level.

##### 3) Lick left-lick right

Next, mice were trained to lick to a specific lick port based on pole location. On each trial, the pole was presented in one of two alternative go locations: the posterior go location or the middle go location. Each pole location was designated a lick port (e.g., posterior with right lick port and middle with left). On presentation of the pole, the mouse was rewarded if it licked the designated lick port. The mouse was punished by timeout if it either licked the non-designated lick port or failed to lick.

##### 4) Lick left-lick right-no lick

Finally, mice were trained on the complete task (‘full task’), involving three pole locations (posterior, middle and anterior) and three behavioural responses (lick left, lick right and do not lick). Once performance reached ∼70% correct, all whiskers except one (C1 or C2) were trimmed to the level of the fur (with retrimming as necessary).

On each protocol, from Go – go no onwards, mice were first trained with trials in blocks of the same type (‘On policy’) and subsequently with trials in a pseudo-random sequence (‘AB policy’):

##### On policy

Here, trials were presented in blocks of the same pole location. The pole location was changed only once the mouse performed a criterion number of consecutive trials (typically 3-8) correctly.

##### Antibias (AB) policy

Here, the type of each trial was determined randomly, subject to the constraint that runs of the same pole location were limited to a maximum (typically 3). We either used the same probability for each trial type (most sessions) or the same probability for go and no go trials. During early training, probabilities could be adjusted in order to correct mouse bias.

During a typical training session in the full task, a few trials at the beginning of the session were delivered using the On policy before switching to the AB policy (Fig. 3-1).

**Figure 3.**
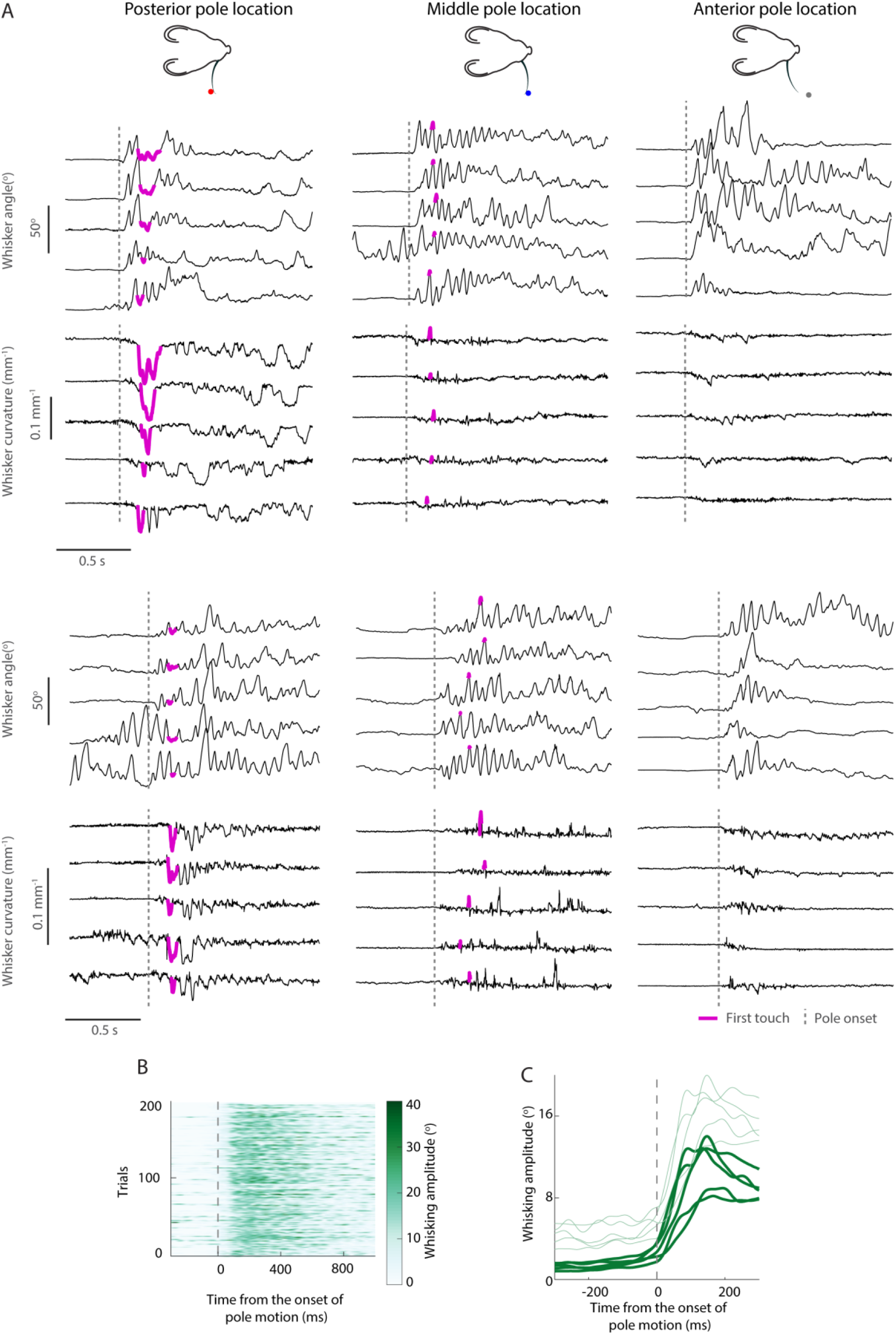
Whisking kinematics and bending during the task. **A.** Example trajectories of whisker angle and whisker curvature for posterior (left panels), middle (middle panels) and anterior (right panels) pole locations in two mice (top and bottom panels). **B.** Whisking amplitude in 200 whisker tracked trials (*Materials and Methods* and Fig. 3-1 and 3-2) for an example mouse, relative to onset of pole movement (vertical broken line). **C.** Mean (thick line) + SD (thin line) whisking amplitude across whisker tracked trials of each mouse. Whisking amplitude significantly increased after pole onset (200ms interval before and after pole onset; t-test p = 4·10^-4^).

### High-speed whisker imaging

Whiskers were imaged as described in O’Connor et al., 2010a and Campagner et al., 2016. Briefly, whiskers ipsilateral to the pole were illuminated from below using an infrared (940 nm) LED array: infrared illumination was used to avoid visual cues to pole location. Whiskers were imaged in the horizontal plane using a high-speed camera (1000 frames/s, 0.4 ms exposure time, Mikrotron).

### Whisker tracking and touch detection

The large number of trials and sessions imaged necessitated automatic whisker tracking requiring minimal user intervention. In this study, we only tracked those sessions in which the mice performed the task at criterion with a single whisker (ca 10^7^ frames, see Fig. 3-1 and 3-2). To extract whisker position/shape from the high-speed imaging data, we first applied the ‘Whisk’ whisker tracker (Clack et al., 2012). The tracker output was then checked by an automated quality-control program to identify misclassified or poorly tracked video frames, based on expected whisker length and location within the image.

To avoid whisker tracking errors close to the face due to fur and whisker pad movement, we used, following Pammer et al. (2013), a face-fur mask. The mask was the mouse snout contour (Bale et al 2015) translated 30 pixels away from the snout border. Whisker bending (curvature) and whisker position (whisker angle) were computed at the intersection of the whisker and the mask by fitting a quadratic curve to a segment of the tracked whisker distal to the mask. Whisker angle was defined as the angle of the tangent to the whisker (at the intersection) with respect to the anterior-posterior axis of the mouse (0°corresponded to the anterior-posterior axis in the nose to tail direction).

In order to detect the onset and offset times of whisker-pole contact with millisecond accuracy, we developed a semi-automatic touch detection GUI (Graphical User Interface). Pole location in each video frame was determined by convolution with a circular pole template. The minimum distance between pole centre and tracked whisker was calculated in each frame and putative touches identified as when this distance was lower than a user-defined threshold. The user then used the GUI to confirm putative touches and to curate their timing to frame-rate precision. In this way, we identified ‘touch episodes’ on each trial, where each touch episode was a continuous sequence of frames, each having a confirmed touch. On each trial, the first touch was classified as protraction or retraction based on the phase of the Hilbert transform of the whisker angle time series (Kleinfeld and Deschênes, 2011) and manual curation. During touch curation, whisker tracking output was also visually inspected. For a subset of the data (5·10^5^ frames), we detected and classified as protraction or retraction all touches in every trial.

If a frame failed the above quality-control procedure, that frame was classified as ‘dropped’. If dropped frames occurred during the first touch, that trial was either re-tracked or discarded. Curvature/angle of occasional, isolated dropped frames was corrected by interpolation of values from adjacent frames.

### Behavioural and imaging data analysis

#### Quantification of learning time and asymptotic performance of mice

In this study, mouse performance (‘task performance’) was quantified as the proportion of trials on which mouse choice was correct during a session. We considered only AB trials of single whisker sessions in which the mouse was performing the full task, and compared the actual performance to that expected if the mouse responded randomly. To this end, we shuffled the pole location sequence with respect to the mouse choice sequence and computed the proportion of correct trials. By repeating this procedure 10000 times, we estimated the mean and 95% confidence interval on task performance attributable to chance. We considered a mouse to have learned the task when performance exceeded the 95% chance confidence interval on three consecutive sessions. We defined asymptotic performance as the performance averaged over eight consecutive, above-chance sessions as close as possible to the end of training (mice 32, 36 and 38) or just before second whisker trimming (mice 33 and 34; Fig. 2 A and 2-1 A).

#### Analysis of whisker movement

In order to quantify whisker movement during the task, we computed whisking amplitude from whisker angle as detailed in Campagner et al. (2016).

#### Classifiers: input and output variables

In order to quantify how well a set of one or more ‘predictor variables’ (sensory variables such as bending moment magnitude and variables reflecting choices on previous trials) might predict a mouse’s choices on a trial and to quantify how much information they contain about the actual pole location, we used a classifier-based approach. Classifiers were trained to predict mouse choice or pole location based on one or more predictor variables obtained from (1) the whisker tracking and touch scoring procedures detailed above and (2) the mouse’s choice on the previous trial. The predictor variables were:

-*Presence/absence of touch*: a binary variable scoring whether or not the whisker touched the pole on a given trial before the mouse choice.

-*Touch type*: a three-valued variable scoring whether the first whisker-pole touch on a trial occurred during retraction or protraction; or, alternatively, if touch was absent.

*-Δκ*_*95*_: a continuous-valued variable measuring bending moment during the first whisker-object touch on a given trial. During touch, a whisker bends. The curvature (*κ*) at a given point along the whisker shaft is equal to the sum of the intrinsic curvature of the unbent whisker and a change in curvature (*Δκ*) due to the whisker-object contact (Solomon and Hartmann, 2006). *Δκ*, at a given point along the whisker shaft, is proportional to the bending moment around the axis normal to the imaging plane through that point (Birdwell et al., 2007; Campagner et al., 2017). *Δκ*_*95*_ is a noise-robust, scalar index of the largest *Δκ* during the first touch episode of a given trial. For each frame *f* of the first touch episode, *Δκ(f)* was computed by subtracting from *κ(f)* the median curvature in the 6 ms before touch onset. The 5^th^ and 95^th^ percentiles of these *Δκ* values were calculated and *Δκ*_*95*_ set equal to whichever had greater absolute value. If no touch occurred during the trial, *Δκ*_*95*_ was, by definition, zero.

*Choice type*: A six-valued variable indicating both the mouse’s choice in a given trial and whether or not it was correct.

#### Classifiers: training and testing procedure

The classifiers used were: PAT classifier (predictor variable was presence/absence of touch), touch type classifier (predictor variable: touch type), *Δκ*_*95*_ classifier (predictor variables: touch type and *Δκ*_*95*_) and previous choice classifier (predictor variable: choice type in the previous trial).

To attempt to classify pole location from predictor variables, we used Maximum a Posteriori (MAP) probabilistic classifiers (implemented in Matlab using the function *fitcnb*). For each mouse, the training/testing data consisted of a vector *Y* specifying the pole location (*y*) on each trial and a matrix *X* specifying the predictor variables on each trial. *Y* consisted of *T* rows: each element *y* was a ternary scalar (k=1,2,3 corresponding to anterior, middle or posterior locations respectively). *X* consisted of *T* rows and *R* columns: each row specified the value of *R* predictor variables (*x*_*1*_,*x*_*2*_,*…,x*_*R*_) on a given trial.

As detailed below, we used the training data to estimate, for each trial, *P*(*y =* k|*x*_1_,…, *x*_*R*_) – the posterior probability that pole location was class *k*, given the predictors:

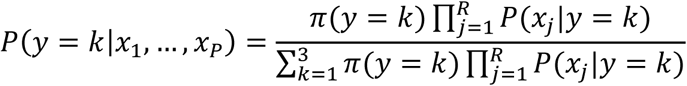

Here *π(y)* is the prior probability of pole location *y* (determined from relative frequencies within the training set) and *P(x*_*j*_*|y)* is the probability of predictor *x*_*j*_ conditional on pole location. The *R* predictors were assumed to be conditionally independent given pole location. For each trial, the pole location predicted by the classifier was set to that with the maximal posterior probability over *k*.

The distributions *P(x*_*j*_*|y)* for categorical predictors were described by multinomials; those for continuous predictors were approximated as Gaussians. Classifier accuracy did not change when the latter distributions were described non-parametrically.

To avoid overfitting, we used 10-fold cross-validation. The trials were randomly allocated across folds. The trials of each fold (10% of the dataset) were used for testing the classifier, with the remainder (90% of the dataset) used for training. Classifier performance was computed after concatenating the prediction outcomes obtained from each of the 10 folds. Classifier chance level, and confidence interval on it, were computed by shuffling the relationship between trial type and mouse choice, and repeating the cross-validation procedure (50 iterations).

We used two different metrics to quantify classification performance. ‘Classifier performance’ was the proportion of trials for which the classifier correctly predicted pole location. ‘Classifier mouse-choice consistency’ was the proportion of trials for which the classifier made the same choice as the mouse (using the mapping between pole location and correct choice defined above).

We also trained classifiers to predict mouse choice instead of pole location. The procedure was as described above, except that *y* specified mouse choice on each trial (a ternary scalar representing whether the response was lick left, lick right or no lick).

#### Quantification of perseveration

Probability of perseveration was computed as the proportion of whisker tracked trials in which choice in the current trial was identical to that in the previous trial. Chance levels for probability of perseveration were computed by random shuffling as described above.

## RESULTS

### The three-choice object localisation task

To investigate active perceptual decision making, our aim was to develop an active touch task which challenges mice to use rich mechanosensory cues whilst allowing the sensory input that guides decisions to be precisely measured trial by trial with millisecond resolution. To this end, we trained mice to perform a novel, three-choice object localisation task with their whiskers (Fig. 1 A-C). Head-fixed animals were trained to use one whisker to localise a metal pole in a dark, sound-proofed enclosure under infra-red illumination. On any given trial, the pole was presented in one of three locations (anterior, middle or posterior) along the anterior-posterior axis of the mouse. Mice were trained to associate each pole location with a unique response: lick at left lick-port (‘left lick’), lick at right lick-port (‘right lick’) or refrain from licking (‘no lick’; for 2 mice the contingencies were reversed, *Materials and Methods*). There were, therefore, nine possible trial-choice outcomes, three correct and six incorrect (Fig. 1 B). For clarity, in the rest of the paper, we label each choice according to the pole location for which that choice was correct. For the example in Fig. 1 B, when the pole was presented in the posterior location, the correct choice was right lick - ‘posterior choice’.

Mice were first trained to perform the task with all whiskers. The number of whiskers was progressively reduced by trimming until, in the final phase of training (‘full task’), mice performed the task with only one whisker (Fig. 2 A and 2-1 A, dark purple dots). Mice learned the full task in 36 ± 12 days of training (mean ± SD across mice) and performed 179 ± 39 trials per daily session (grand mean across both mice and sessions ± SD of session-means across mice; Fig. 2 A and 2-1 B). We expressed a mouse’s ‘task performance’ as the proportion of trials on which its choice was correct. Mice reached stable task performance of 0.74 ± 0.08 (grand mean ± SD of session-means; *Materials and Methods*) and the performance of all mice was above chance (Fig. 2 B). To verify that mice were relying on their whiskers to perform the task, we trimmed the whisker of fully trained mice and retested. As expected, task performance dropped significantly (t-test; p = 0.0013; Fig. 2 C-D) from 0.72 ± 0.04 (grand mean ± SD of session-means) pre-trim to 0.36 ± 0.02 post-trim. Post-trim performance was within 95% confidence interval of chance (Fig 2 C-D; *Materials and Methods*). In sum, these results indicate that mice can learn a three-choice object localisation task using a single whisker.

### High-speed imaging and whisker tracking

The fact that mice localised the pole using only one whisker drastically limits the sensory input available to the mouse to guide its decisions, and makes it feasible to experimentally measure that input on a trial-by-trial basis. To investigate how mice made choices on the task, we used high-speed imaging (1000 frames/s) both to measure whisker movement and to estimate whisker bending during whisker-pole touch (Fig. 1 A right panel and 1 C). Due to the high volume of imaging data (∼3×10^8^ frames), we selected for detailed analysis 7.4 ± 2.7 sessions per mouse, where the animal was performing the full task with a single whisker. For analyses of these data, we pooled trials across sessions: thus task performance is reported as mean ± SD across mice: the data comprised 761 ± 175 trials per mouse (Fig. 3-1 and 3-2). Task performance in these sessions (0.74 ± 0.05) was consistent with that reported above and was above chance at all three pole locations (posterior 0.77 ± 0.07; middle 0.75 ± 0.06; anterior 0.70 ± 0.06; Fig. 4 A).

**Figure 4.**
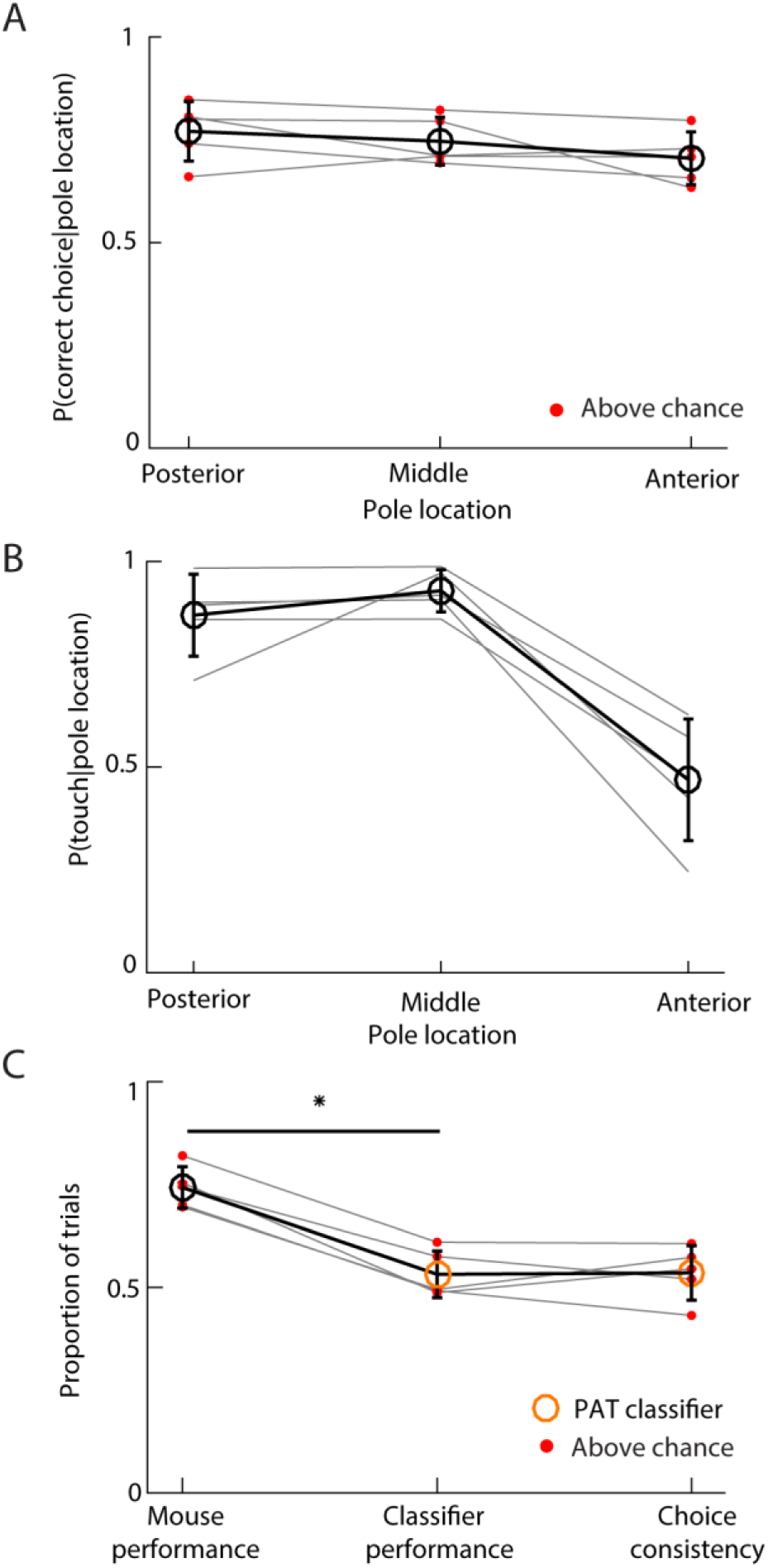
Presence/absence of touch (PAT) cannot account for mouse choice. **A.** Probability of correct choice as a function of pole location, for each mouse (gray lines). Empty circles and error bars show mean and SD across mice. Red dots indicate that performance of a given mouse was outside 95% confidence interval on chance (5000 shufflings). **B.** Probability of touch as a function of pole location. Gray lines indicate individual mice. Empty circles and error bars: mean and SD across mice. **C.** Task performance of mouse and PAT classifier along with choice consistency. Red dots indicate that the classifier/mouse performance or choice consistency was significantly higher than chance. Empty circles: mean. Error bars: SD. *: t-test p =1.8·10^-4^.

We tracked the location and shape of the whisker in every frame of the selected sessions (*Materials and Methods*). To quantify whisker movement (‘kinematics’), we extracted the angle of the whisker near its base. As a proxy for bending moment, we measured the curvature of the whisker near its base relative to its intrinsic, contact-free value (Fig. 1 A right panel, 1 C, 3 A and B).

Consistent with previous work on two-choice pole localisation (O’Connor et al., 2010a; Guo et al., 2014b), we found that mice adopted a stereotyped whisking strategy. At the start of a trial, prior to pole movement, mice whisked little (Fig. 1 A right panel, 1 C, 3 A and 3 B). Shortly after the onset of pole movement, all mice started to whisk (Fig. 3 C).

### Whisker bending direction and magnitude predict mouse choice

To investigate the mechanosensory cues that informed mouse choices, we first applied a touch detection algorithm to the imaging data to register, on each trial, whether or not a mouse touched the pole with its whisker (*Materials and Methods*), and tested whether the most elementary cue, presence/absence of touch on a given trial (PAT), might be informative. We found that touches occurred at all pole locations: almost always at both posterior (0.87 ± 0.10) and middle (0.93 ± 0.05) locations; less often (0.46 ± 0.15; t-tests p<0.004) at the anterior location (mean ± SD across mice; Fig 4 B). This suggests that PAT is unlikely to fully differentiate pole location. To test this quantitatively, we computed the ability of a probabilistic classifier (‘PAT classifier’) to predict pole location from PAT only (*Materials and Methods*). We measured classifier performance, in the same way as mouse performance, as the proportion of trials for which it predicted pole location correctly. We measured classifier-mouse choice consistency (abbreviated to ‘choice consistency’) as the fraction of trials in which mouse and classifier made the same choice. We found that performance of the PAT classifier (0.53 ± 0.06; mean ± SD across mice) was significantly lower than that of the mice (0.74 ± 0.05; t-test p = 1.8·10^-4^) and that choice consistency was mediocre (0.53 ± 0.07; Fig. 4 C), but above chance (red dots in Fig. 4 C, right column; *Materials and Methods*). These results confirm that this task challenges mice to use sensory cues richer than PAT.

Which additional mechanosensory cues might be guiding mouse choice? Primary Whisker Neurons (PWNs) are sensitive to the direction of whisker deflection (Gibson and Welker, 1983a; Lichtenstein et al., 1990; Bale and Petersen, 2009; Bale et al., 2013; Maravall et al., 2013) and have recently been shown to encode both the direction and magnitude of the bending moment associated with whisker-object active contact during behaviour (Campagner et al., 2016; Severson et al., 2017). We wondered whether these cues – information that is redundant in simpler tasks – might account for the mice performance.

To test whether bending moment direction might be an informative cue, we first classified each trial according to whether the first whisker-pole touch on the trial occurred during protraction or retraction. Retraction and protraction touches cause bending in opposite directions (Fig. 3 A). The ‘touch type’ on each trial was scored from the imaging data as either ‘no touch’, ‘protraction touch’ or ‘retraction touch’ (Fig. 5 A). First touch was a good proxy for subsequent touches on a given trial. 84% of trials had at most three touches; 94% of second touches were of identical type to the first and 98% of third touches of identical type to the second.

**Figure 5.**
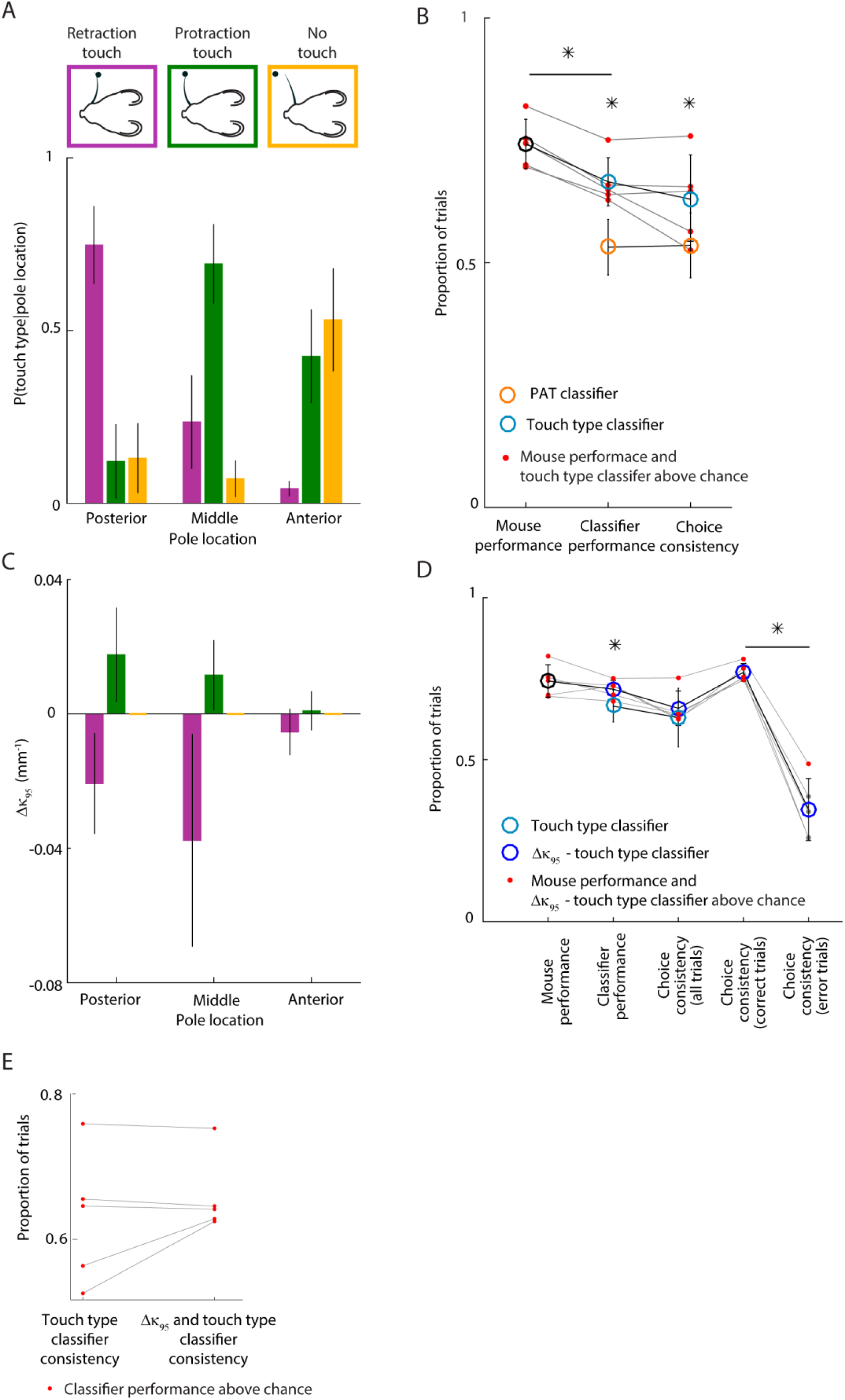
Whisker bending magnitude and direction account for mouse choice. **A.** Probability of each touch type as a function of pole location: mean and SD across mice. **B.** Performance of touch-based classifiers compared to mice. Red dots indicate that corresponding classifier/mouse performance or choice consistency was significantly greater than chance for the given mouse. Orange circles show mean classifier performance and choice consistency of the PAT classifier (same data as Fig. 4 C). Error bars: SD. *: t-test, p ≤ 0.05. **C.** Mean *Δκ*_*95*_ of each touch type as a function of pole location for an example mouse. Error bars: SD across trials. **D.** Performance of touch and bending based classifiers compared to mice. Red dots indicate that corresponding classifier /mouse performance or choice consistency was significantly greater than chance for that mouse. Light blue show mean classifier performance and choice consistency of the touch type classifier (same data as Fig. 5 B). Error bars: SD. *: t-test, p≤0.05. **E.** Single mouse values of touch type only and touch type - Δκ_95_ choice consistency.

We found that touch types differed in frequency at each pole location (Fig. 5 A; one-way ANOVAs, p <10^-5^). The posterior pole location tended to elicit retraction touch; the middle location protraction touch, and the anterior location no touch or protraction touch. This suggests that mouse whisking strategy was to adjust whisking set point to a position intermediate between the middle and posterior pole locations. In this way, whisking would tend to cause whisker-pole contact during protraction for the anterior/middle locations and contact during retraction for the posterior location. These data indicate that direction of touch could potentially be a useful cue. To test this, we used the classifier approach to quantify how well pole location on a trial could be predicted from touch type (‘touch type classifier’). We found that the touch type classifier not only performed better than the PAT classifier (0.67 ±0.05 vs 0.53 ± 0.06; t-test, p = 0.0011), but also that its choice consistency was higher (0.63 ± 0.09 vs 0.53 ± 0.07; t-test p = 0.0066; Fig. 5 B), although to variable extent across mice (Fig 5 E). However, the touch type classifier performed significantly worse than the mice (0.74 ±0.05; t-test, p = 5.5·10^-4^). Thus, touch type is informative, but not sufficient, to account fully for mouse performance.

We considered the possibility that mice might be able to use a continuous readout of bending moment as a cue. When a whisker strikes an object, it bends and its curvature changes. We computed a simple index sensitive to bending moment magnitude during first touch, termed *Δκ*_*95*_ (*Materials and Methods*). *Δκ*_*95*_ is a robust measure of the most extreme value of curvature change (*Δκ*) during a given touch. *Δκ*_*95*_ was variable, but depended systematically on pole location (Fig. 3 A and 5 C). Protraction touch was typically associated with positive *Δκ*_*95*_ (one tailed t-test, p = 7·10^-5^), retraction touch with negative *Δκ*_*95*_ (p = 0.0043). For each mouse, for retraction and protraction touches, magnitude of *Δκ*_*95*_ was dependent on pole location (two-way ANOVAs, p<10-^8^). These data suggest that bending moment magnitude is a potential cue to pole location. To test whether *Δκ*_*95*_ might permit improved task performance compared to touch type alone, we again used the classifier approach. We trained a classifier given input of both *Δκ*_*95*_ and touch type to predict pole location (*Materials and Methods*). This classifier performed as well as the mice (0.72 ± 0.03 vs 0.74 ± 0.05 respectively; t-test, p =0.21) and, overall, significantly better than the touch type classifier (Fig. 5 D; t-test, p = 0.038). Choice consistency for the *Δκ*_*95*_ classifier was 0.66 ± 0.05 (mean ± SD across mice), but variable across mice (Fig. 5 E). For the three mice where consistency between mouse choice and the touch type classifier was highest, the *Δκ*_*95*_ classifier failed to increase choice consistency. In contrast, for the two mice where consistency between mouse choice and touch type classifier was lowest, the *Δκ*_*95*_ classifier increased choice consistency. These findings indicate that bending moment strength and direction – quantities that PWNs are known to encode – can account for the ability of mice to perform the task substantially more accurately than a strategy based purely on presence/absence of touch. The findings also indicate that individual mice differ in the exact weight that different mechanical variables have in their decisions.

### Choices on previous trials predict performance on error trials

The analysis above considered both trials where the mouse chose correctly (‘correct trials’) and those where it chose incorrectly (‘error trials’). To get further insight into mouse decision making, we selectively investigated errors (Fig. 1 B). One possibility is that errors might be driven by current sensory input – for example, due to an unusual touch on a particular trial. Alternatively, errors might be driven by memory of outcomes on previous trials (Kiyonaga et al., 2017; Akrami et al., 2018). We asked how well the best of the classifiers considered above (that with both touch type and *Δκ*_*95*_ as inputs) could predict mouse choice on error trials. Consistent with the data reported above, this classifier was accurate on correct trials (0.77 ± 0.03). In contrast, the classifier was remarkably inaccurate on error trials: on average, choice consistency on error trials was significantly lower than that on correct trials (0.35 ± 0.1 vs 0.77 ± 0.03; paired t-test p = 4·10^-4^) and for only one mouse was it above chance (Fig. 5 D). This suggests that there might be an important non-sensory contribution to choices on error trials.

To test for a possible contribution to choice from previous trial outcomes, we first examined the time sequence of mouse choices. We found that mice showed a strong tendency to make the same choice on consecutive trials; that is, to perseverate (Fig. 6 A - B). The probability of perseveration on error trials (0.63 ± 0.02) was substantially above chance for all individual mice and significantly greater than that on correct trials (0.63 ± 0.02 vs 0.36 ± 0.02; paired t-test, p=1.4·10^-5^). Perseveration was not simply a consequence of response bias since those mice for which the three choice types were statistically equally likely (χ^2^ test, p>0.26), still showed significant perseveration. The most common perseverating behaviour leading to error, was that, when a mouse got a trial correct, it tended to repeat the successful choice on the next trial (Fig. 6 D). Indeed, when tested on error trials, a classifier trained to predict choice based on that in the previous trial was substantially more accurate than a classifier trained to predict choice based on sensory input (0.58 ± 0.08 vs 0.35 ± 0.1; t-test p = 0.0076; Fig 6 C). In contrast, when tested on correct trials, the choice-based classifier was less accurate (0.41 ± 0.03 vs 0.77 ± 0.04; t-test p = 4·10^-5^; Fig 6 C). Taken together, these results indicate that two competing mechanisms governed mouse decision-making during the task, driven by choice-memory and current sensory input respectively.

**Figure 6.**
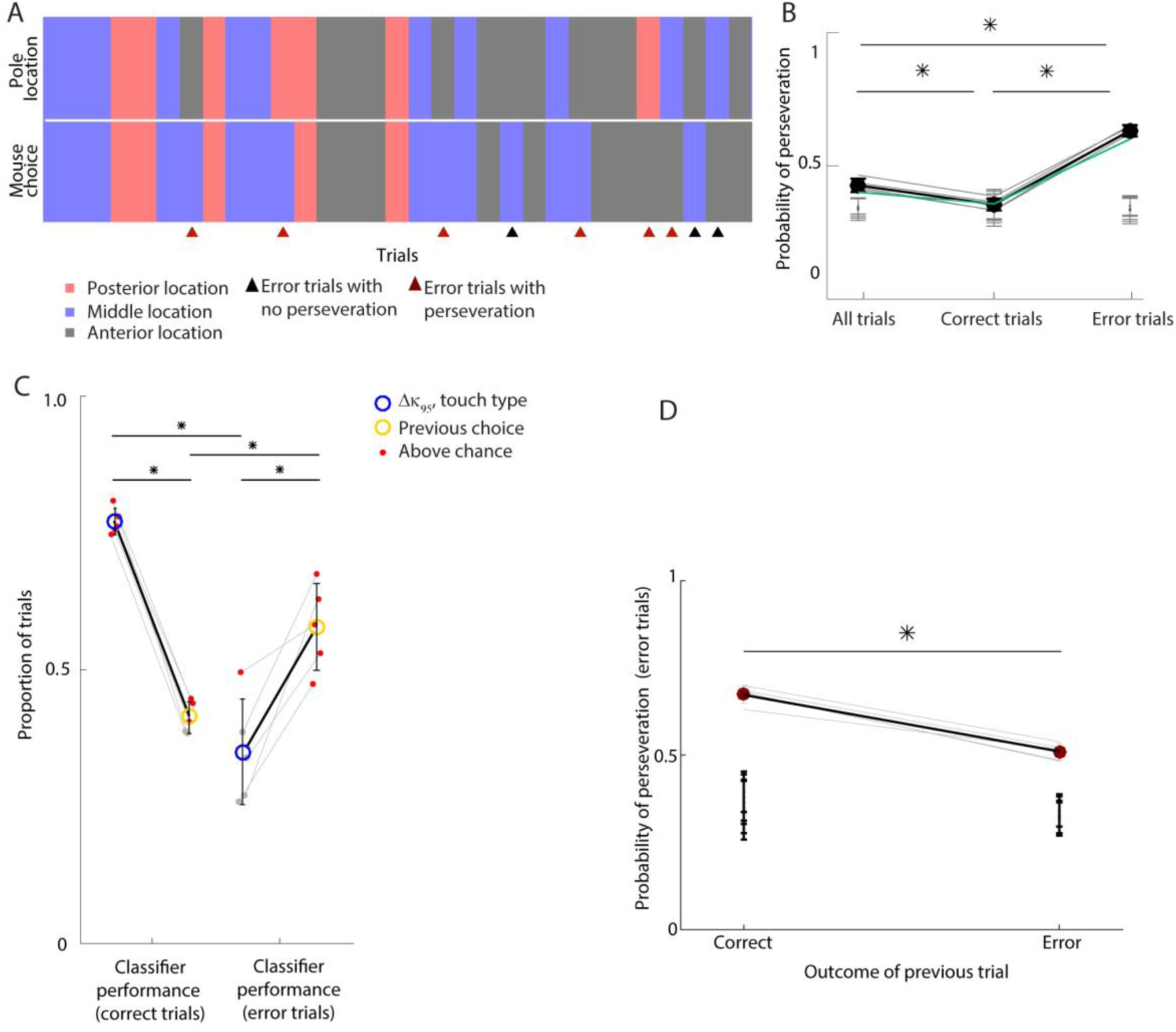
Previous choice predicts error trials. **A.** Sequence of 31 consecutive trials performed by an example mouse. Red, blue and gray rectangles indicate trials in which the pole location was anterior, middle or posterior (the top row), or in which the mouse made a posterior, middle or anterior choice (bottom row). Triangles indicate error trials, dark red triangle indicate error trials in which the choices in the previous and current trial were identical (i.e. the mouse perseverated). **B.** Probability of perseveration for each mouse (gray lines) under different conditions: considering all trials (left), correct trials only (middle) or error trials only (right). Dark green line indicates example mouse in panel A. Black circles: mean, black error bar: SD across mice. *: t-test, p ≤ 0.0167 (Bonferroni correction, n = 3). Gray bars indicate chance interval (10000 shuffling, 95% confidence interval). **C.** Performance of classifiers predicting mouse choice correct trials only (left) and error trials only (right). Blue and yellow circles indicate mean values for Δκ_95_ - touch type classifier, and previous choice classifier respectively. Small dots are single mice values. Red indicates that the classifier performance value for the mouse was above chance. Error bar are SD across mice. *: t-test, p ≤ 0.05. See Fig. 6-1 B. **D.** Probability of perseveration during error trials depending on whether the previous trial was a correct trial or an error trial. *: t-test, p = 3.4·10^-4^. Black error bars indicate chance intervals of each mouse (10000 shufflings, 95% confidence interval).

## DISCUSSION

When making perceptual decisions under natural conditions, animals move their sense organs (‘active sensation’). We developed a new active sensation task which challenges mice to use multiple mechanosensory cues, whilst allowing the sensory input that drives decisions to be measured at millisecond resolution. In this three-choice task, mice use a single whisker to localise a pole. We found that competing sensory and internal processes influenced decision making, and identified both mechanosensory and choice-memory signals that accurately predicted mouse choice.

### A new task for investigation of active perceptual decision making

Our study builds on previous work which developed whisker-based object localisation in head-fixed mice, along with a mechanics framework and experimental methods for estimating the mechanical forces associated with whisker-pole interaction (Birdwell et al., 2007; O’Connor et al., 2010a; Clack et al., 2012; Pammer et al., 2013; Campagner et al., 2016, 2017). Our task is novel compared to previous rodent object localisation tasks in that it is a three-choice task. The task maintains the ability to estimate whisker mechanical forces, but requires animals to use multiple mechanosensory cues, including the direction of bending moment.

### Mechanosensory basis of active touch

We found that correct choices could be predicted with high accuracy from the direction and magnitude of whisker bending. Neurons throughout the whisker system are sensitive to the direction of passive whisker deflection (Gibson and Welker, 1983a; Simons and Carvell, 1989; Lichtenstein et al., 1990; Bale and Petersen, 2009; Maravall et al., 2013). During active whisker-object contact, the activity of PWNs primarily reflects bending moment: torque generated as contraction of the whisking muscles cause the whiskers to bend against the object (Campagner et al., 2016; Severson et al., 2017; Bush et al., 2016; reviewed by Campagner et al., 2017). PWNs robustly encode both the direction and magnitude of bending and transmit this information along the ascending thalamo-cortical pathway (Yu et al., 2006, 2016; O’Connor et al., 2010b; Huber et al., 2012; Petreanu et al., 2012; Xu et al., 2012; Hires et al., 2015; Moore et al., 2015; Peron et al., 2015b; Gutnisky et al., 2017). A wide range of PWN properties (Zucker and Welker, 1969; Gibson and Welker, 1983b; Lichtenstein et al., 1990; Szwed et al., 2003; Jones et al., 2004; Arabzadeh et al., 2005; Leiser and Moxon, 2007; Bale and Petersen, 2009; Lottem and Azouz, 2011; Bale et al., 2013; Maravall et al., 2013) can be concisely explained by this framework (Campagner et al., 2017). Thus, the cues we found to predict choices are consistent with physiological properties of somatosensory neurons. They are also consistent with biomechanical modelling studies (Yang and Hartmann, 2016; Huet et al., 2017).

Sensing of bending moment provides a simple account for how rodents solve a number of whisker-dependent tasks. Mice solve two-choice, anterior-posterior pole localisation tasks by a selective whisking strategy. The strength and number of touches is sufficient to guide to pole location (*Introduction;* O’Connor et al., 2010a). In our three-choice task, mice whisked in such a way that they contacted the pole at all three locations. Mice solved the task by focussing their whisking at a location intermediate between the anterior and posterior pole locations. In this way, on trials where the pole was located anterior/middle, touch typically occurred during the forward (protraction) phase of whisking whereas, on trials where the pole was posterior, touch typically occurred during the backward (retraction) phase. Thus, direction of bending was informative about pole location. In addition, touches at the anterior location, when they occurred at all, were weaker (bending magnitude was lower) than those at the posterior/middle locations, so that bending magnitude was also informative about pole location. In addition to object localisation, sensing of bending moment also accounts for wall-following behaviours (Sofroniew et al., 2014). Sensing of bending moment may also permit whisker-based inference of object shape (Solomon and Hartmann, 2006) and of the spatial structure of the environment (Fox et al., 2012; Pearson et al., 2013). Some active touch tasks may require multidimensional mechanosensory signals – for example, axial force in combination with bending moment (Bagdasarian et al., 2013; Pammer et al., 2013). The role of bending moment in texture discrimination tasks, which have mainly been analysed in terms of stick-slip events (Wolfe et al., 2008), requires further research: dynamic signals, such as rate of change of bending moment, may be important here. Overall, bending moment sensing provides both a paradigm for future investigation of neural algorithms of active touch and an inspiration for further development of tactile robotics.

### Competing contributions to perceptual decision-making from sensory input and choice-memory

We found that correct choices were predicted from immediate sensory information with no detectable effect of previous choices, whereas incorrect choices were predicted from previous choice with no detectable effect of immediate sensory information. Choice-history-dependence is consistent with previous studies of other sensory systems, but has not previously been reported in the tactile domain (Busse et al., 2011; Fassihi et al., 2014b; Marcos and Harvey, 2016; Hwang et al., 2017; Kiyonaga et al., 2017; Akrami et al., 2018). This double dissociation suggests two distinct neural systems competing to influence decisions: one driven by immediate sensory information; the other driven by memory of previous choices. Although a choice-memory-guided system might improve performance in a task where the sequence of trials is predictable, when, as in our task, the sequence is random, history-dependence leads to errors, whilst correct choices necessarily depend entirely on immediate sensory information (Kiyonaga et al., 2017; Akrami et al., 2018). The sensory-guided system is likely to involve the ascending sensory pathway through the primary somatosensory cortex (S1). S1 neurons respond robustly to both magnitude and direction of whisker bending (O’Connor et al., 2010b; Hires et al., 2015; Peron et al., 2015b; Yu et al., 2016; Kwon et al., 2017; Martini et al., 2017) and inactivation of S1 impedes correct choices on active whisking tasks, including pole localisation (O’Connor et al., 2010a; Guo et al., 2014a) and wall following (Sofroniew et al., 2015). The choice-memory-guided system may involve a widely distributed circuit (Hanks and Summerfield, 2017), with recent research pointing to a particular role for posterior parietal cortex (Raposo et al., 2014; Akrami et al., 2018).

In summary, we have developed a new, tactile object localisation task that permits high resolution measurement of the mechanosensory input that drives perceptual decisions. The task has shed new light both on the mechanical mechanisms of active touch and on how sensory input and choice-memory interact to influence decisions. In future studies, the task can be combined with cellular-resolution measurement of neural activity, and may serve as a useful tool for investigating how competing sensory and internal neural mechanisms contribute to active perceptual decision making.

## EXTENDED DATA

**Figure 2-1.**
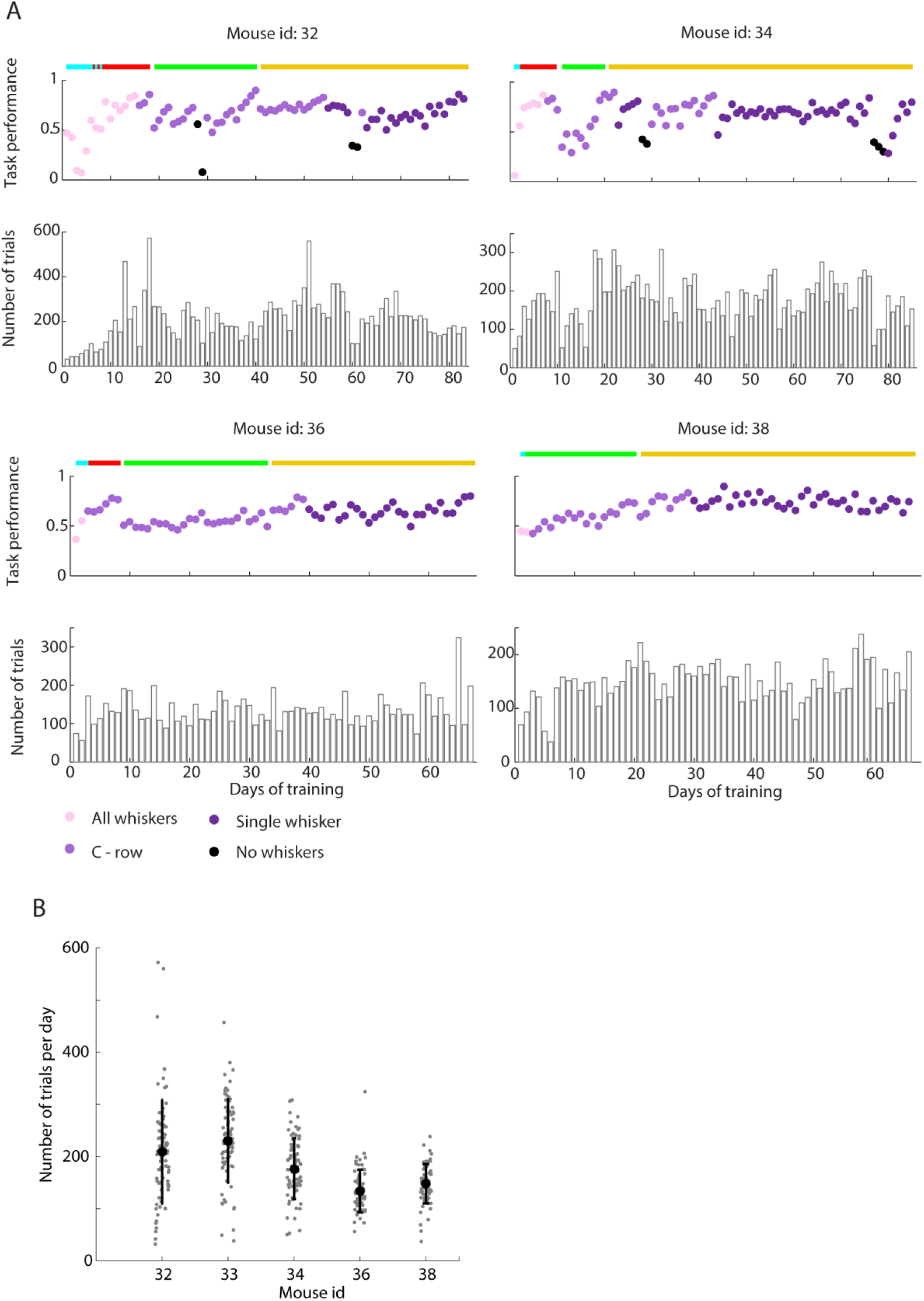
The three-choice object localization task. Data from mice not shown in Fig. 2 A and total number of trials for each mouse. A. Learning curves and total number of trials performed each day of those mice not shown in figure 2 A. Note that in two mice (36 and 38), whiskers were trimmed to C row at the conclusion of lick protocol (cyan bar). In one mouse (38) go - no go protocol (red bar) was skipped. After the mouse learnt the lick protocol, we immediately introduced the lick left-lick right protocol (green bar). The no go location was introduced with ‘the full task’ (gold bar). B. Total number of trials per day performed by each mouse (black circles: mean, error bars: SD).

**Figure 3-1.**
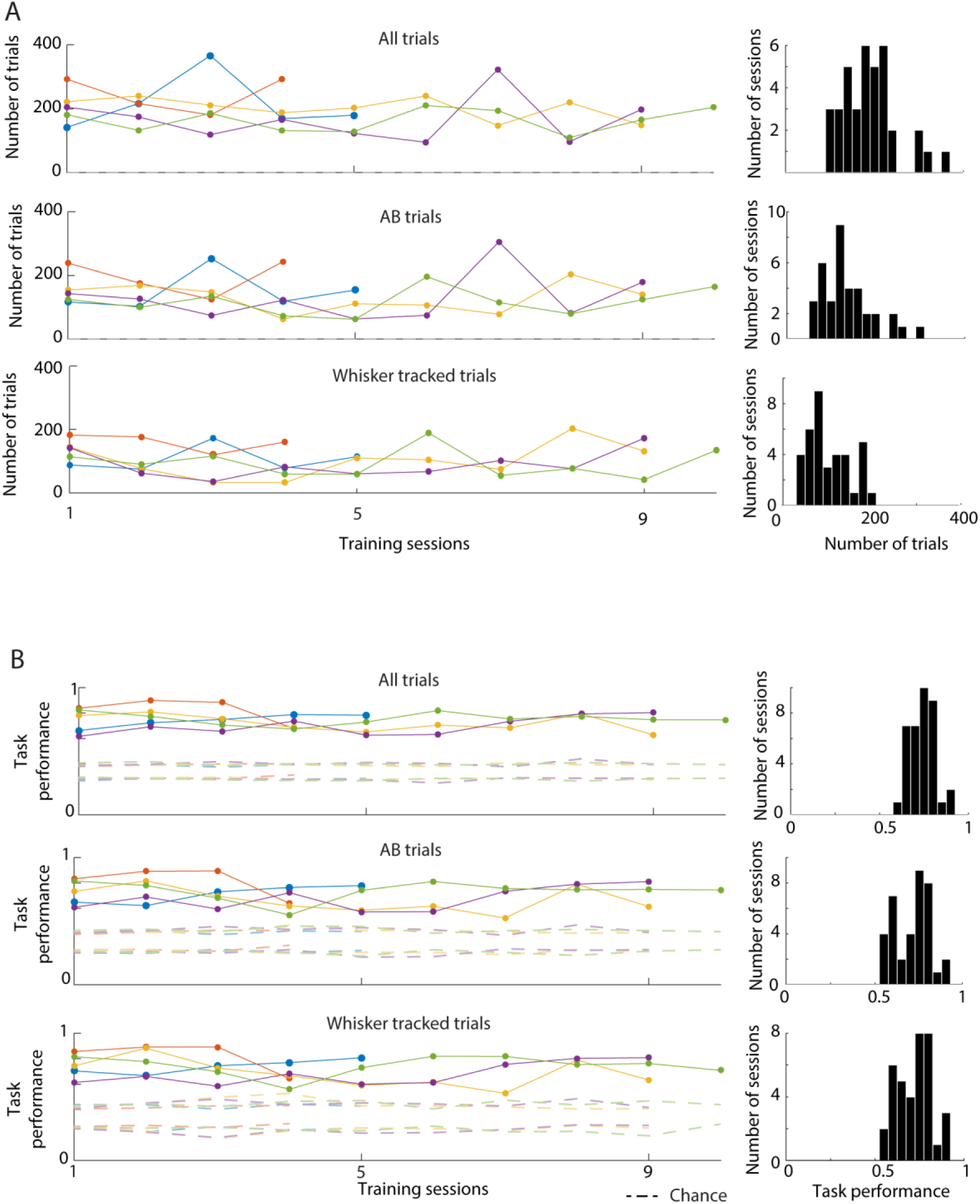
Mouse performance during the whisker tracked sessions. **A. Left:** number of trials in the sessions where the whisker was tracked. Dashed lines represent the bounds of the 95% chance confidence interval for each mouse and session (computed as for figure 2 B; 10000 shufflings). **Right**: histogram of number of trials shown in the respective left panel, pooling together the data of all mice. In both A and B different colours indicate different mice. **B. Left**: task performance of the sessions shown in **A.** Dashed lines represent the bounds of the 95% chance confidence interval for each mouse and session (10000 shufflings). **Right**: histogram of performances shown in the respective left panel, pooling together the data of all mice.

**Figure 3-2.**
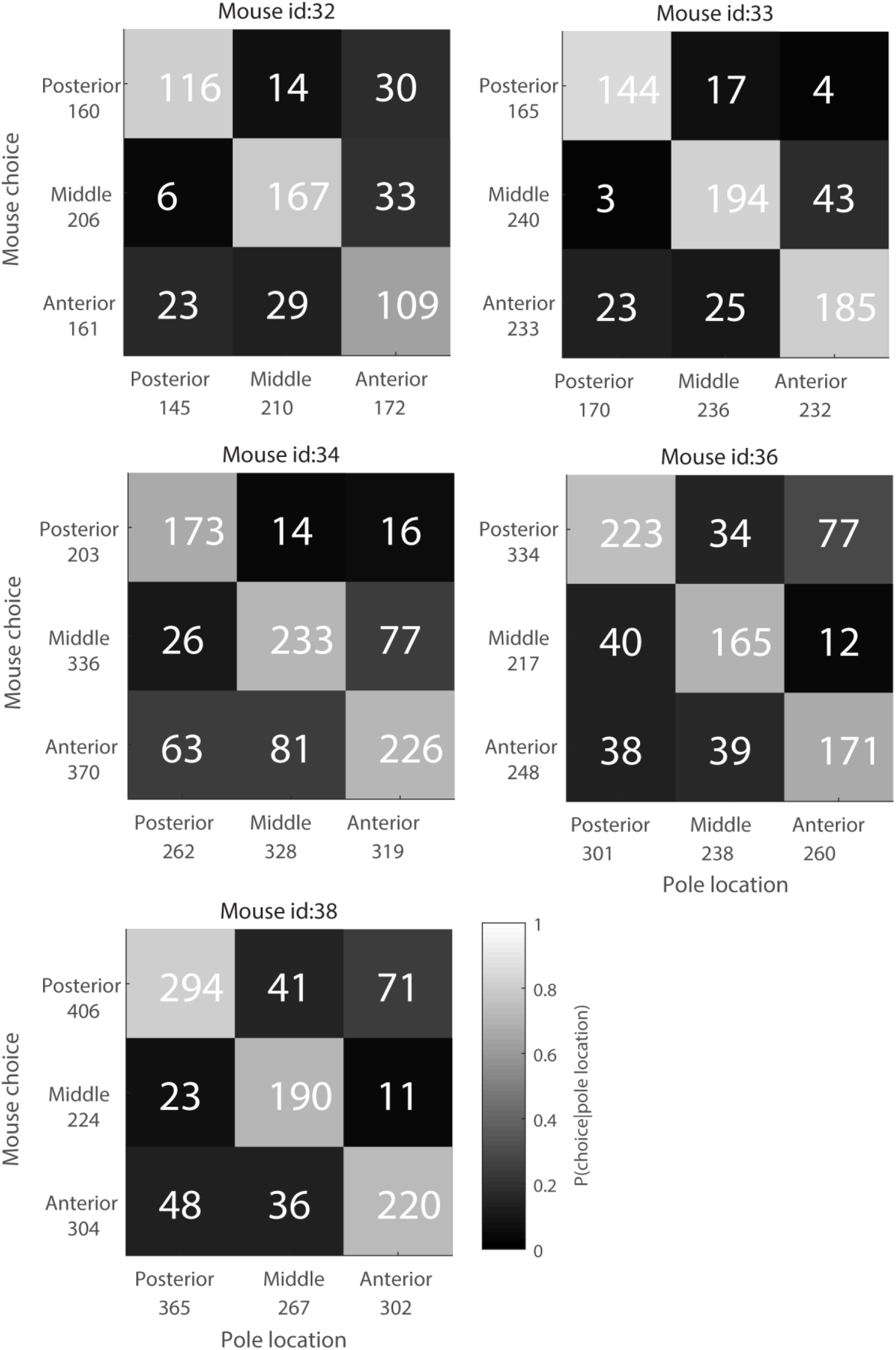
Mouse choice at each pole location for whisker tracked trials. Confusion matrices summarizing the choices made by each mouse, at each pole location, using data only from whisker tracked trials. Each matrix shows (white numbers inside the matrix) the number of trials on which each of the 9 possible location-choice possibilities occurred and (gray shading) the conditional probability of each possible choice given each possible pole location. The total number of trials for each choice and each pole location are also shown (left and below matrix respectively).

Author contribution
Designed research: CD, RSP, KS
Performed research: CD, KC, DP, SF
Contributed unpublished analytic tools: MHE, MH
Analyzed data: CD, MHE,ACR, MSEL, SF
Wrote the first draft of the paper: CD, RSP
Edited the paper: CD, RSP, MHE, KS, MH

## Acknowledgements

We dedicate this paper to the memory of Katarina Chlebikova. We thank Luciana Walendy and Nuo Li for their support with behavioural training; Robert Lucas for comments on the manuscript; Tiago Branco and Branco laboratory members for discussion. This work was funded by Biotechnology and Biological Sciences Research Council (BB/L007282/1, BB/ P021603/1), Medical Research Council (MR/L01064X7/1, MR/P005659/1), Wellcome Trust (097820/Z/11/B) and HHMI.

